# Quantitative variation within a species for traits underpinning C_4_ photosynthesis

**DOI:** 10.1101/253211

**Authors:** Gregory Reeves, Pallavi Singh, Timo A. Rossberg, E. O. Deedi Sogbohossou, M. Eric Schranz, Julian M. Hibberd

## Abstract

Engineering C_4_ photosynthesis into C_3_ crops such as rice or wheat could substantially increase their yield by alleviating photorespiratory losses^1,2^. This objective is challenging because the C_4_ pathway involves complex modifications to the biochemistry, cell biology and anatomy of leaves^3^. Forward genetics has provided limited insight into the mechanistic basis of these characteristics and there have been no reports of significant quantitative intra-specific variation of C_4_ attributes that would allow trait-mapping^4,5^. Here we show that accessions of C_4_ *Gynandropsis gynandra* collected from locations across Africa and Asia exhibit natural variation in key characteristics of C_4_ photosynthesis. Variable traits include bundle sheath size and vein density, gas exchange parameters and carbon-isotope discrimination associated with the C_4_ state, but also abundance of transcripts encoding core enzymes of the C_4_ cycle. Traits relating to water use showed more quantitative variation than those associated with carbon assimilation. We propose variation in these traits likely adapted the hydraulic system for increased water use efficiency rather than improving carbon fixation, indicating that selection pressure may drive C_4_ diversity in *G. gynandra* by acting to modify water use rather than photosynthesis. As these accessions can be easily crossed and produce fertile offspring, our findings indicate that natural variation within a C_4_ species is sufficiently large to allow genetic-mapping of key anatomical C_4_ traits and regulators.

---

Plants that use C_4_ photosynthesis can effectively abolish photorespiratory losses caused when Ribulose 1,5-Bisphosphate Carboxylase/Oxygenase (RuBisCO) fixes oxygen rather than CO_2_ ^6,7^. In C_4_ plants, RuBisCO is typically sequestered in bundle sheath (BS) cells that are concentrically arranged around the vasculature. Establishment of a molecular CO_2_ pump delivers carbon to RuBisCO from Mesophyll (M) cells via C_4_ acid intermediates^8^. C_4_ photosynthesis relies on an increased importance of the BS for photosynthesis, reduced dependence on M cells, more chloroplasts in BS cells, increased proliferation of plasmodesmata between M and BS cells, and a higher vein density to increase the volume of the leaf occupied by the BS. These morphological alterations to the leaf that facilitate the C_4_ cycle are known as Kranz anatomy^9^. Moreover, photosynthesis gene expression is modified such that genes encoding components of the C_4_ and Calvin-Benson-Bassham cycles are strongly and preferentially expressed in either M or BS cells^8,10^.

Despite the complex modifications associated with C_4_ photosynthesis, current estimates are that the C_4_ pathway has evolved independently more than sixty times in angiosperms^11^, which suggests a relatively straightforward route must allow the transition from the ancestral C_3_ to the derived C_4_ state. Genome-wide analysis of transcript abundance in multiple C_3_ and C_4_ species has provided unbiased insight into processes that likely change in C_4_ compared with C_3_ leaves^12–14^. Furthermore, *cis-*elements that control expression of genes encoding the C_4_ cycle have been documented. To date however, the regulators that recognize these motifs have not been isolated^15^. Despite progress in our understanding of C_4_ photosynthesis, it is currently not possible to rationally design a C_4_ pathway in a C_3_ leaf.

When natural variation is present, it enables quantitative methods such as Genome-Wide Association Studies (GWAS) and/or the development of a mapping population. Molecular marker-trait associations on the population allow identification of the causal genes underpinning the variation, which has been used extensively to map loci responsible for numerous complex traits in plants^16^. If such an approach could be applied to study C_4_ photosynthesis, then it would expedite discovery of key regulators to engineer increased photosynthetic efficiency in C_3_ plants. Interspecific hybrids have been generated between C_3_ and C_4_ species of the dicotyledon *Atriplex*^17^. Although progeny possessed variation in C_4_ phenotypes, specific traits showed limited penetrance and there were high rates of sterility^18^. In the grasses, *Alloteropsis semialata* shows natural variation in C_4_ parameters and has been classified into C_3_ or C_4_ subspecies^19,20^, but there are currently no reports that these populations have been bred. Thus, to our knowledge there are currently no examples that variation in C_4_ traits within a single species is sufficient to allow breeding and then molecular trait-mapping. We therefore investigated the extent to which key C_4_ traits varied in the C_4_ dicotyledonous *Gynandropsis gynandra,* which is a leafy green vegetable^21^ in a clade with both C_3_ and C_4_ species^22–24^. Here we show that accessions of *G. gynandra* show significant variation in both anatomical and physiological aspects associated with C_4_ photosynthesis. These accessions have short generation spans, are sexually compatible and produce fertile offspring. These findings indicate that in a dicotyledonous species that is phylogenetically close to the model *Arabidopsis thaliana* there is sufficient natural variation to allow the use of classical genetics to identify loci controlling the multifaceted C_4_ syndrome.

Accessions of *G. gynandra* were collected from African and Asian sub-continents were used (Supplementary Table 1). DNA sequencing and phylogenetic reconstruction generated a taxonomy that was generally consistent with geographical origin but also indicated that the accession from Benin in West Africa was more like the Asian accessions than those from East Africa (Fig. 1a). These accessions displayed considerable variation in macroscopic characters associated with leaf appearance (Fig. 1b, Supplementary Fig.1a). For example, fully expanded leaflets varied in size and shape, and there was also variation in petiole length, presence of trichomes and anthocyanin pigmentation. As there was considerable macroscopic variation in leaf characteristics, we then evaluated these accessions for variation in features of Kranz anatomy. Interestingly, there were statistically significant differences in vein density (Fig. 1c&d, Supplementary Fig.1b), cross-sectional area of BS strands (Fig. 1c&e, Supplementary Fig. 1c), size of individual BS cells (Fig. 1f) and stomatal density (Fig. 1c&g). Furthermore, East African accessions showed higher vein density, reduced distance between veins, and a greater stomatal density than Asian lines (Supplementary Fig. 3a-c). Asian accessions typically had larger BS areas and cell sizes than those from the African continent (Supplementary Fig. 3d&e). Vein density was inversely correlated with BS area and BS cell size but positively with stomatal density (Supplementary Table 2). The average number of BS cells around each vein showed no statistically significant differences between lines (Supplementary Fig. 3f), but cross-sectional area of the BS and the size of individual BS cells were positively correlated (*ρ*=0.8, *P*<0.0001). We therefore conclude that the area of individual BS cells, rather than the number of these cells per vein bundle, drives the increased BS strand area. This suggests that genetic determinants of cell size rather than cell proliferation are involved in the variation in BS tissue in *G. gynandra*. Thus, despite the lower phenotypic variation associated with C_4_ compared with C_3_ leaves^25^, our findings demonstrate flexibility is still possible within individual species that are fully C_4_.

**Figure 1.**
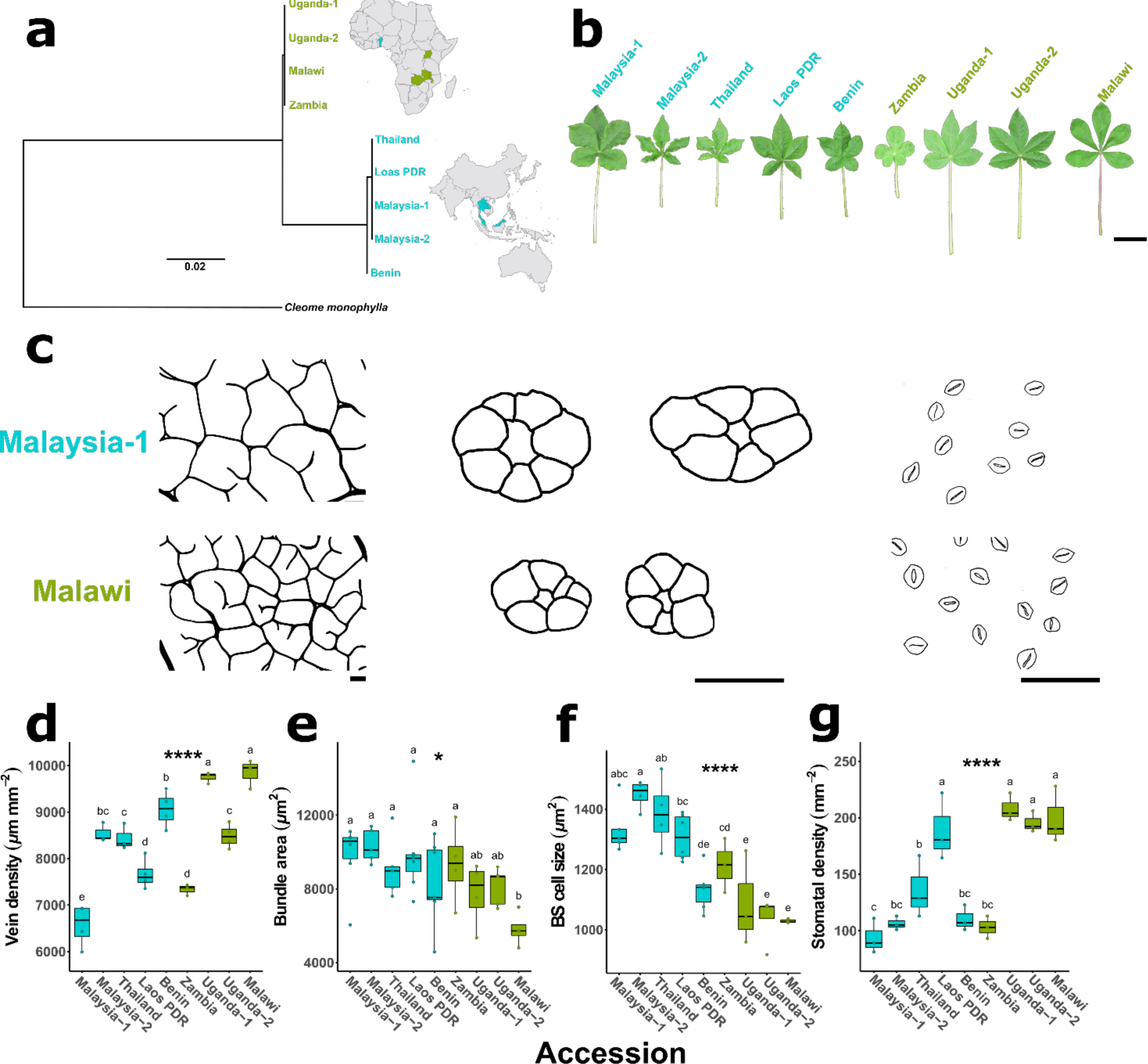
Natural variation in Kranz anatomy features among a diverse panel of *G. gynandra*(C_4_) accessions. **a**, Geographic and phylogenetic relationships for nine accessions from seven countries across Africa and Asia. **b,** variation in fully mature whole leaves of six-week old plants (scale = 5 cm). **c**, variation in venation, bundle sheath ring, bundle sheath cell size, and stomata traces of fully mature leaves for two extreme examples (scale = 100 µm). **d**, vein density, **e**, bundle area, **f**, bundle sheath cell size, **g**, and stomata density for all accessions. Asterisks indicate significant differences between accessions (one-way ANOVA, \**P*<0.05, \*\**P*<0.01, \*\*\**P*<0.001, \*\*\*\**P*<0.0001). Letters above individual box-scatter plots indicate significant groupings according to Duncan’s Multiple Range Test (*α*=0.05).

We next investigated whether differences observed in Kranz anatomy affected photosynthetic performance. For all accessions, their CO_2_ response curves (assimilation (*A*) response to the concentration of CO_2_ inside the leaf *C*_*i*_) were typical of C_4_ plants with high carboxylation efficiencies and low CO_2_ compensation points *Γ* (Fig. 2a, Supplementary Fig. 4a). Although parameters associated with instantaneous gas exchange such as maximum rate of photosynthesis (*A*_*max*_), rate of photosynthesis under the conditions of growth (*A*_*400*_), CO_2_ carboxylation efficiencies and *Γ* showed little variation between accessions (Fig. 2b-e, Supplementary Fig. 4b-e), there were statistically significant differences in transpiration (Fig. 2f), stomatal conductance (Fig. 2g) and water use efficiency *WUE* (Fig. 2h). Furthermore, there was also significant variation in the carbon isotope discrimination against ^13^C (*δ*^*13*^*C*) in leaf dry matter (Fig. 2i), which is a measure of the efficiency of the C_4_ carbon pump over the life-time of the leaf. Asian accessions showed reduced discrimination against *δ*^*13*^*C* compared with East African lines (Supplementary Fig. 4i). These data therefore indicate that the accessions of *G. gynandra* possess significant variation in parameters linked to the balance between water use and photosynthesis that influenced the efficiency of the C_4_ cycle over the lifetime of a leaf.

**Figure 2.**
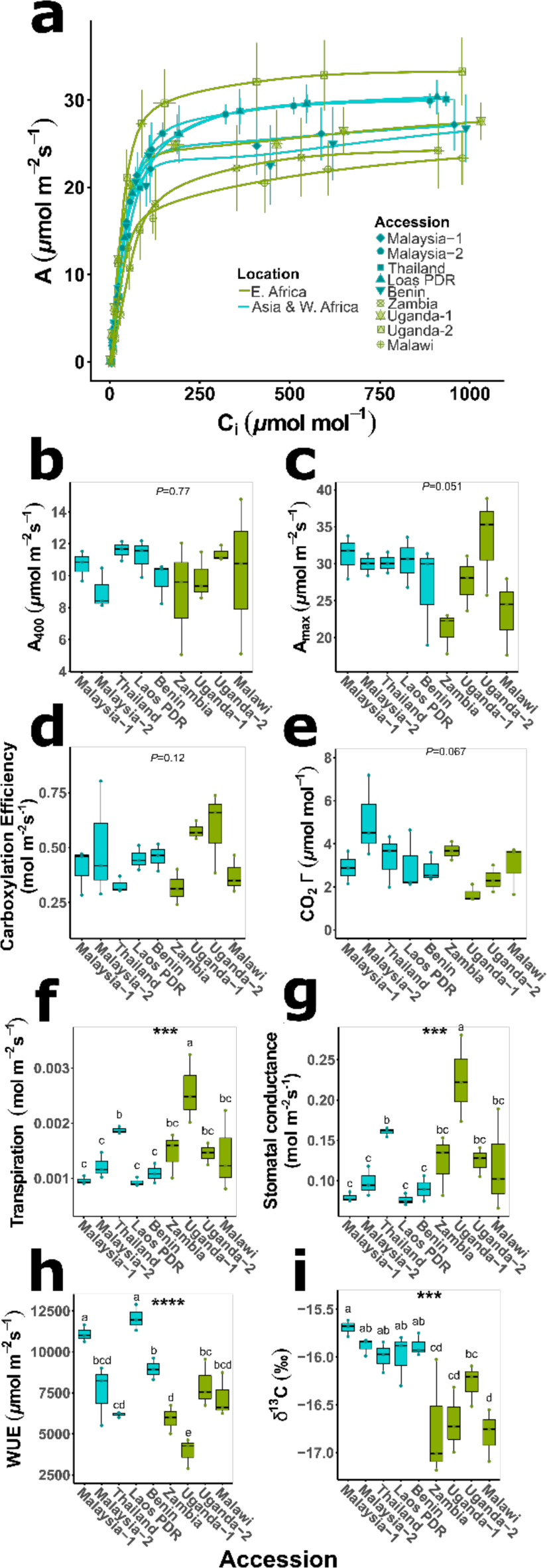
Physiological variation for photosynthetic gas exchange parameters among a diverse panel of *G. gynandra* (C_4_) accessions. **a**, assimilation (*A*) verses internal CO_2_ (*C*_*i*_) response curve. **b-i**, differences among accessions for ambient assimilation (*A*_*400*_) rates (400 ppm atmospheric [CO_2_], *C*_*a*_; PPFD 350 µmol m^-2^s^-1^), maximal assimilation (*A*_*max*_) rates (1200ppm *C*_*a*_, PPFD 2000 µmol m^-2^s^-1^), CO_2_ compensation point (*Γ*), carboxylation efficiency, transpiration, stomatal conductance, water use efficiency (*WUE*), and carbon isotope composition (*δ*^*13*^*C*), respectively. Asterisks indicate significant differences between accessions (one-way ANOVA, \**P*<0.05, \*\**P*<0.01, \*\*\**P*<0.001, \*\*\*\**P*<0.0001). Letters above individual box-scatter plots indicate significant groupings according to Duncan’s Multiple Range Test (*α*=0.05), n=3.

We next sought to investigate the extent to which transcript abundance of core genes of the C_4_ cycle differed between the accessions. Interestingly, there were statistically significant differences in the abundance of transcripts encoding Phospho*enol*pyruvate carboxylase (*PEPC*) which catalyses the first committed step of the C_4_ cycle, the BS-specific decarboxylase NAD-dependent Malic Enzyme (*NAD-ME*), the small subunit of RuBisCO (*RbcS*), and pyruvate,orthophosphate dikinase (*PPDK*) that regenerates PEP the primary acceptor of 
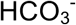
 (Fig. 3a,c,e,g). In all cases, these differences in C_4_ transcript abundance were associated with geographical location and phylogenetic position of the accessions, with Asian and West African accessions accumulating greater levels of C_4_ transcripts than East African accessions (Fig. 3b,d,f,h). Understanding how photosynthesis enzymes become strongly expressed and patterned to either mesophyll or bundle sheath cells of C_4_ species is a longstanding area of research. However, although progress has been made in understanding *cis*-elements responsible, there is little known about the transcription factors involved. The intraspecific variation in expression of genes encoding enzymes of the C_4_ cycle in *G. gynandra* therefore provides an opportunity to identify *trans*-factors important for C_4_ photosynthesis.

**Figure 3.**
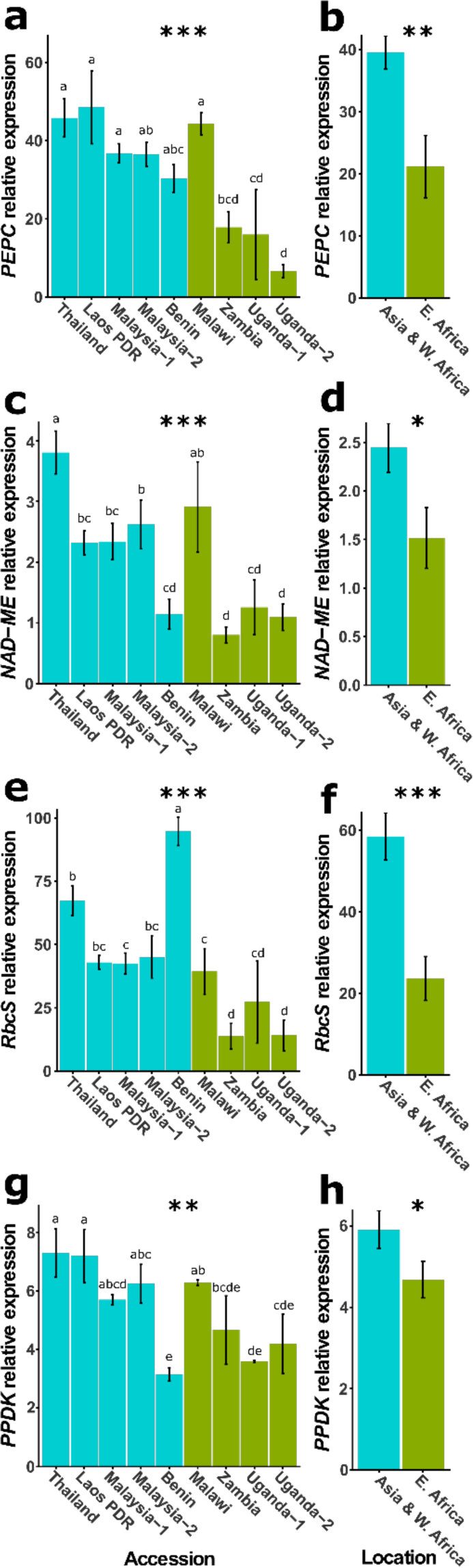
Transcript abundance differences for key enzymes in the C_4_cycle among diverse *G.gynandra* accessions. Gene expression differences were determined by qRT-PCR. Geneabbreviations: *PEPC*, *PHOSPHO*ENOL*PYRUVATE CARBOXYLASE 2*; *NAD-ME*, *NAD-DEPENDENT MALIC ENZYME 2*; *RbcS*, *RIBULOSE-1,5-BISPHOSPHATE CARBOXYLASE/OXYGENASE SMALL SUBUNIT 1A*; *PPDK*, *PYRUVATE,ORTHOPHOSPHATE DIKINASE*. Asterisks indicate significant differences (\**P*<0.05, \*\**P*<0.01, \*\*\**P*<0.001, \*\*\*\**P*<0.0001), **a**, **c**, **e**, **g**, among accessions (one-way ANOVA, n=3) or **b**, **d**, **f**, **h**, among phylogenetic cluster (Student’s t-test, n=15 for Asia and W. Africa, n=12 for E. Africa). Letters above individual bar charts indicate significant groupings among accessions according to Duncan’s Multiple Range Test (*α*=0.05), n=3.

Despite accessions functioning with similar photosynthetic efficiencies under ambient CO_2_ and light conditions, when assessed by phylogenetic grouping those with more pronounced Kranz traits (*e.g*., larger BS tissues and lower vein densities) exhibited increased *A*_*max*_, *WUE* and *δ*^*13*^*C* (Supplementary Fig. 4c,h,i) and stronger expression of the C_4_ cycle (Fig. 3b,d,f,h). To summarize, compared with East African accessions, Asian and West African accessions tended to have higher *WUE*, lower density of stomata and veins, and larger BS areas and cell sizes. Lastly, consistent with the higher *δ*^13^*C*, which is indicative of a stronger C_4_ cycle, the Asian and West African accessions had increased expression of genes encoding C_4_ enzymes (Fig. 4).

**Figure 4.**
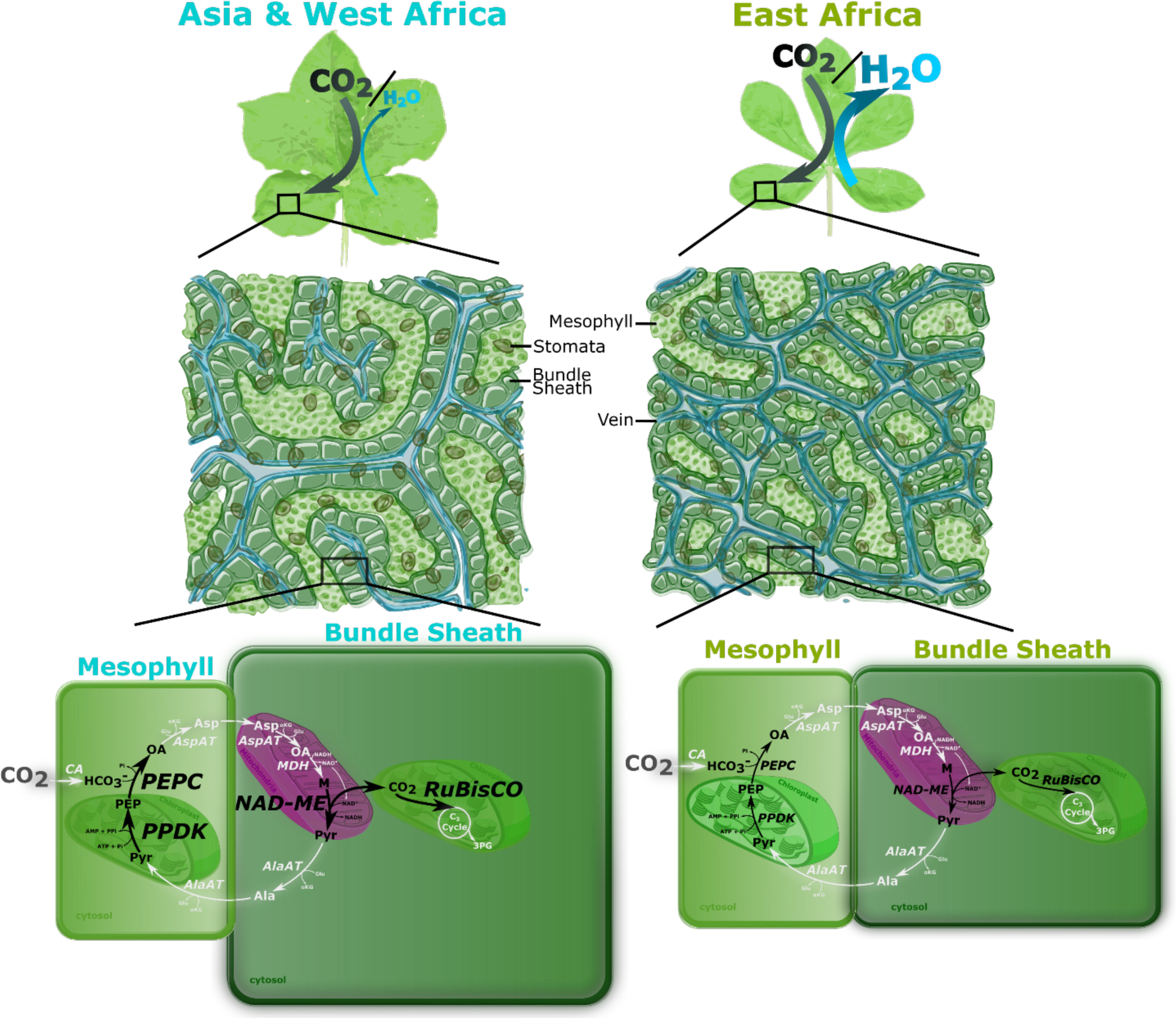
Asian and African *G. gynandra* accessions exhibit many differences in anatomy, physiology and C_4_ enzyme expression patterns. All C_4_ enzymes investigated had differential transcript abundance and are indicated by black arrows, where larger letters represent higher relative transcript abundance. Gene abbreviations: *PEPC*, *PHOSPHO*ENOL*PYRUVATECARBOXYLASE 2*; *NAD-ME*, *NAD-DEPENDENT MALIC ENZYME 2*; *RbcS*, *RIBULOSE-1,5-BISPHOSPHATE CARBOXYLASE/OXYGENASE SMALL SUBUNIT 1A*; *PPDK*, *PYRUVATE,ORTHOPHOSPHATE DIKINASE.* Enzymatic steps in white were not investigated.

The considerable variation reported in this study offers a valuable germplasm resource to identify regulators of the C_4_ pathway and Kranz anatomy through genetic mapping. All accessions in this study hybridize easily. Emasculation and pollination need only take 15-30 seconds per flower. For example, the most divergent accessions regarding anatomy ‘Malaysia-1’ X ‘Malawi’, ‘Malaysia-2’ X ‘Malawi’, and their reciprocal crosses produce an average 52 ± 11 seeds per silique (n=6), whose offspring are fully fertile. These F_1_ hybrid populations provide an excellent breeding foundation to delineate regulatory mechanisms, and also provide an opportunity to test whether the C_4_ trait is induced by a master switch^26^, or the action of multiple independent processes^15,27^. The discovery of intra-specific variation in a C_4_ grass would be particularly useful in mapping traits relevant to improving photosynthesis in cereals and thus introduce C_4_ photosynthesis into C_3_ crops.

While our understanding of the regulatory mechanisms underlying C_4_ metabolism is growing, there is still a significant gap in tools to expand understanding of the regulation behind Kranz anatomy and the C_4_ biochemical cycle. Methods such as Quantitative Trait Loci (QTL) mapping or GWAS in *G. gynandra* or in an equally diverse C_4_ species may provide beneficial insights for the regulation of Kranz development. Most trait variation in *G. gynandra* was associated with characteristics relating to water use that impact on carbon capture. It is noteworthy that modifications to C_3_ leaves considered to represent early steps on the path towards the C_4_ phenotype are also associated with water use rather than CO_2_ fixation^27,28^. As natural vegetation is not considered to be under strong selection pressure to optimize photosynthesis^29,30^, it seems likely that C_4_ trait variation continues to be driven by optimizing water use rather than photosynthesis *per se*.

## ACKNOWLEDGEMENTS

We thank Frank Becker and Beatrice Landoni (Wageningen University) for providing ITS sequence data and initial information on the selected lines, Blandine Gilbert for help with image processing and James Rolfe for carbon isotope analysis. GR was supported by a Gates Cambridge Trust PhD Fellowship and PS by Advanced ERC grant 694733 Revolution to JMH.

## AUTHOR CONTRIBUTIONS

GR, PS and JMH designed the study. GR, PS, TAR and EODS carried out experimental work. GR, PS, TAR, MES and JMH wrote the manuscript.

## MATERIALS AND METHODS

### Plant accessions and growth conditions

A selection of nine diverse accessions of *G. gynandra* were made from a larger germplasm collection based on initial phenotypic and genetic screening. Five accessions were from Africa and four from Asia (Supplementary Table 1, materials available on request from MES). Plants from all *G.gynandra* accessions were grown under identical conditions prior to sampling. After germination, all seeds were planted in 5:1 F2 compost (Levington Advance, UK) to fine vermiculite premixed with 0.17 g/L insecticide (Imidasect 5GR, Fargro, UK). Seedlings were kept in a growth cabinet at 350 µmol photons m^-2^ s^-1^ light with a 16 h photoperiod, at 25 °C, 60% relative humidity (RH), ambient [CO_2_]. A single dose of 3 mL/L slow release 17N-9P-11K fertilizer (All Purpose Continuous Release Plant Food, Miracle-Gro, UK) was applied after 1.5 weeks. Plants for physiological measurements were grown under identical conditions to those for Kranz measurements for the first three weeks, after which they were re-planted in 13 cm^3^ pots with 5:1 M3 soil (Levington Advance, UK) to medium vermiculite soil mixture and moved to a growth room set to 23 °C, 60% RH, ambient [CO_2_], 350 µmol photons m^-2^ s^-1^ PAR with a 16 h photoperiod.

### Preparation of leaf tissue sections

Three weeks after germination, tissue was harvested from healthy plants from the centre trifoliate leaves of the second pair of fully expanded true leaves. A 3 mm^2^ rectangle was cut from the leaf adjacent to the midvein with a razor blade for transverse sections. Two slightly larger rectangles were cut from identical regions for paradermal sectioning and qRT-PCR analysis.

For transverse sections, leaf tissue in plastic cuvettes were submerged in a 4% paraformaldehyde in PBS solution (Sigma-Aldrich, St. Louis, MO, USA), placed in a vacuum chamber for 1 h, and incubated at 4 °C overnight for fixing. Cuvettes then underwent an ethanol dehydration series from 30% to 90% (v/v) ethanol solutions (Thermo Fisher Scientific, Waltham, MA, USA) in 10% (v/v) increments for 45 minutes each at 4 °C with a final overnight treatment at 4 °C in 95% ethanol with 0.1% eosin dye solution (Sigma-Aldrich, St. Louis, MO, USA). The dye solution was washed thrice with 100% (v/v) ethanol at room temperature. The samples were embedded in resin in accordance with the Technovit 7100 (Kulzer GmbH, Wehrheim, Germany) manufacturer’s protocol. Hardened resin blocks were cut with a manual rotary microtome (Thermo Fisher Scientific, Waltham, MA, USA). Sections were placed on microscope slides and stained with 0.1% (w/v) toluidine blue solution (Sigma-Aldrich, St. Louis, MO, USA) prior to imaging on a light microscope.

For paradermal sections, fresh tissue samples were placed in plastic cuvettes and incubated in 3:1 100% (v/v) ethanol to acetic acid solution before treatment with 70% (v/v) ethanol solution (refreshed once) at 37 °C overnight. To clear the samples, cuvettes were submerged in 5% NaOH solution for three hours at 37 °C. After storage in 70% ethanol solution, the samples were stained with 95% (v/v) ethanol and 0.1% (v/v) eosin dye solution (Sigma-Aldrich, St. Louis, MO, USA). Samples were stored overnight at 4 °C and washed with 70% (v/v) ethanol thrice before transfer to slides for imaging. To determine stomatal density impressions of the abaxial epidermis of each central leaflet were generated by applying a thin coat of transparent nail varnish (Boots, Nottingham, UK). After drying, the varnish was peeled off and mounted onto glass slide for imaging.

### Measurement of Kranz anatomy traits

Slides of all leaf sections were imaged with an Olympus BX41 light microscope with a mounted Micropublisher 3.3 RTV camera (Q Imaging, Surrey, BC, Canada). Images were captured with Q-Capture Pro 7 software, and measurements were analyzed with the software ImageJ^31^. To maximize comparability, strict criteria were applied for all image analyses. Microscopy of transverse leaf sections was used to quantify the BS both in terms of average BS tissue area (the total cross-sectional area of all BS cells immediately surrounding a vein) and BS cell size (the average cross-sectional area of individual BS cells around the vein). To quantify BS tissue area, the freehand selection tool was used to subtract the integrated area of each vein from the integrated area of all BS cells in direct contact with the vein on images with 200X total magnification. This value was divided by the number of BS cells in each vein bundle to obtain the average BS cell size. For intervein distance (the distance between the centers of adjacent veins in transverse sections), only vein bundles were measured for which the following criteria did not apply: wide (indicates branching) or extremely large veins, veins with distorted BS cells due to contact with adjacent BS tissues (indicates merging), veins with damaged BS cells. The line selection tool was used to measure the linear distance between the centers of adjacent veins on images with 40X total magnification. Vein density (vein length per unit area of leaf) was quantified on paradermal sections on images with 100X total magnification. Slides were imaged with the same microscopy equipment as transverse sections but set to Ph3 (phase contrast). Three images (from three different leaves per plant) were randomly selected for measurement. The freehand line tool was used to trace all veins (both major and minor) along their center. As it was not possible to trace all veins in an image simultaneously, individual vein sections were progressively measured without overlap and the individual lengths summed. The total vein length was divided by the image area to obtain the density. Stomatal density (the number of stomata per unit area of leaf) was measured on three subsampled leaves from three random plants on images with 200X total magnification. The total number of stomata were divided by the image area to obtain the density.

### Photosynthetic performance

A LI-6800 portable photosynthesis infrared gas analyzer (IRGA) system (LI-COR, Lincoln, NE, USA) equipped with a multiphase flash fluorimeter was used to assess physiological differences for photosynthetic parameters between *G. gynandra* accessions. All physiological measurements were performed on the central leaflet of five-week old plants with three biological replicates. For stomatal conductance, transpiration (*E*), and assimilation (*A*_*400*_), measurements were taken during ambient conditions of growth (400 ppm atmospheric [CO_2_], *C*_*a*_; photosynthetic photon flux density (PPFD) 350 µmol m^-2^s^-1^). Water use efficiency (*WUE*) was defined as *A*_400_/*E*. A combination chlorophyll fluorescence and assimilation / intracellular CO_2_ concentration (*A*/*C*_*i*_) curve was measured for three plants from each accession. Atmospheric CO_2_ (*C*_*a*_) reference values were: 400, 400, 300, 200, 100, 50, 25, 400, 400, 400, 600, 800, 1000, 1200, 400 ppm, with a saturating rectangular pulse of 12,000 µmol m^-2^s^-1^ at each reference point. Otherwise, measurements were made at a PPFD of 2000 µmol m^-2^s^-1^, 23 °C and 60% RH at each reference point. All leaves covered the full area of the fluorimeter/IRGA cuvette. Measurements were carried out on consecutive days between one and eight hours post dawn, measuring one random plant from each accession per day. Maximal assimilation (*A*_*max*_) was calculated as the asymptote of the *A/C_i_* response curve. The CO_2_ compensation point (*Γ*) was calculated from the regression of *A* and *C_i_* measurements ranging between *C*
_*a*_ values of 200 and 25 ppm *at A*= 0. Adjusted *R^2^* values for the regression line ranged between 0.9932 and 0.9967. Carboxylation efficiency was calculated as the partial derivative 
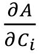
 at *A* = 0. Stable carbon isotope (*δ^13^C*) analysis was performed according to methods previously described^32^ on three biological replicates per accession with 500 µg of dried leaf tissue.

### Statistical Analysis

For all tests, individual plants were considered experimental units in a complete randomized design. Data were analyzed in SAS (University Version, SAS Institute, Cary, NC, USA) and in R (Version 3.4.2, R Studio, Inc., Boston, MA, USA). A One-Way Analysis of Variance (ANOVA) compared all means from anatomical and physiological measurements among *G. gynandra* accessions and a Student’s t-test was used to compare means of accessions by continent (*α*=0.05). Null hypotheses were rejected for specific ANOVA or t-tests for any population with *P* value ≤ 0.05. Levene’s Test was used to evaluate homoscedasticity^33^. Duncan’s Multiple Range post-hoc test was used for mean separation on accessions (*α*=0.05) with statistically significant ANOVAs^34^. Pearson product-moment correlation coefficients were calculated to find associations among features of Kranz traits^35^.

### Analysis of transcript abundance

Leaf tissue samples for RNA extraction were harvested simultaneously with samples for Kranz trait measurements. The fresh samples were immediately frozen with liquid nitrogen and stored at −80 °C. Total RNA was extracted from three tissue samples per accession with a RNeasy Mini Kit (QIAGEN, Hilden, DE) according to the manufacturer’s instructions. An On-Column DNase Digestion protocol was applied to remove genomic DNA contamination (QIAGEN, Hilden, DE) before cDNA was synthesized with Invitrogen Superscript II RT enzyme according to the manufacturer’s instructions (Thermo Fisher Scientific Inc., Waltham, MA, USA). All cDNA samples were stored at −20 °C before qRT-PCR. Primers were designed for Quantitative PCR of C_4_ cycle genes *PEPC, NAD-ME, RbcS* and *PPDK* (Supplementary Table 3), and reactions carried out as reported previously^36^ on three biological and three technical replicates.

## Supplementary Figures

**Supplementary Figure 1.**
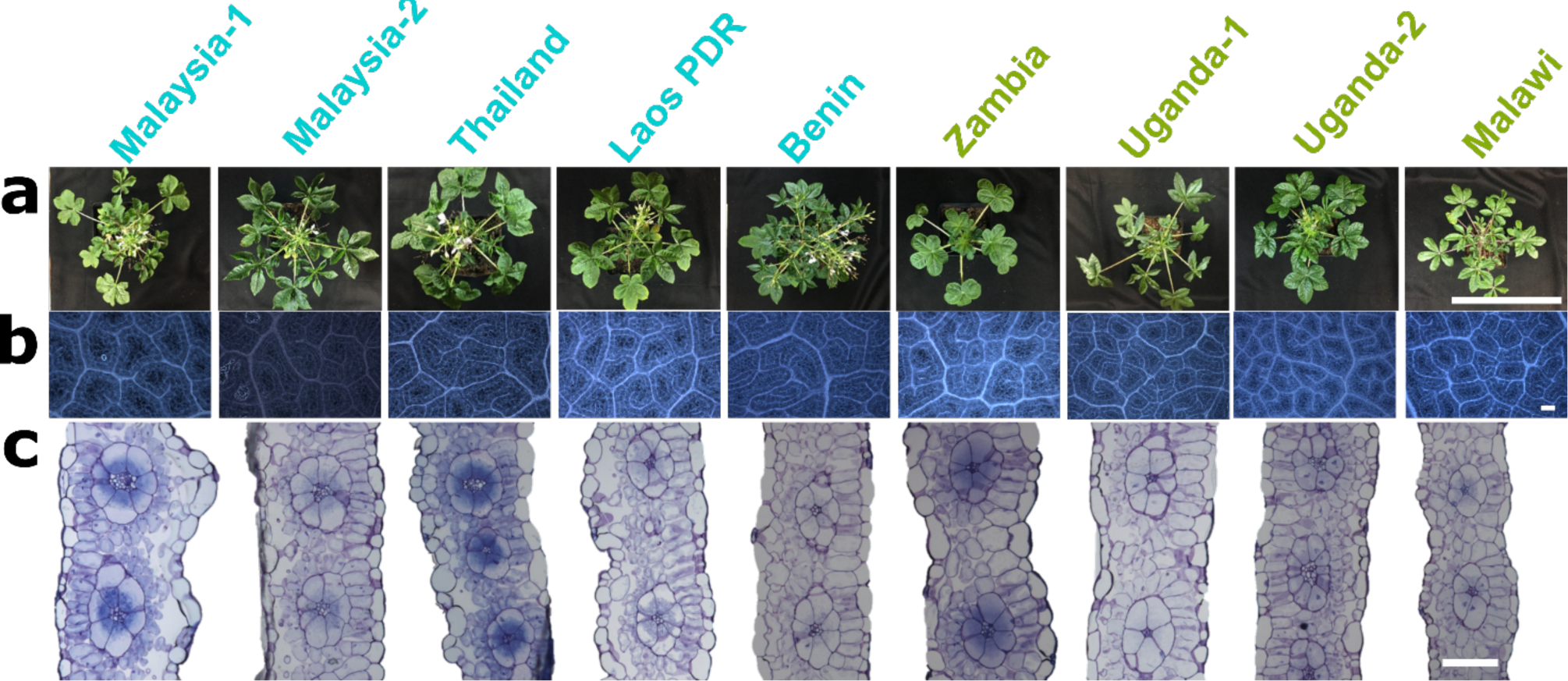
Representative images for macroscopic and microscopic variation in leaf anatomy across a panel of *G. gynandra* accessions. **a**, four-week old whole plants (scale = 30 cm), **b**, paradermal view of leaf venation under light phase contrast microscopy, **c**, transverse leaf sections (scale = 100 µm).

**Supplementary Table 1.** Accessions of *G. gynandra* investigated and their source regions.

| Accession | Temporary No.<sup>1</sup> | VI No.<sup>1</sup> | Continent of Origin | Country of Origin | Source |
| --- | --- | --- | --- | --- | --- |
| Malaysia-1 | TOT7199 | VI055200 | Asia | Malaysia | Prof. Eric Schranz |
| Malaysia-2 | N/A | N/A | Asia | Malaysia | B&T World Seeds<sup>2</sup> |
| Thailand | TOT4937 | VI048669 | Asia | Thailand | Prof. Eric Schranz |
| Laos PDR | TOT7441 | VI055576 | Asia | Lao People's Democratic Republic | Prof. Eric Schranz |
| Benin | ODS-15-020 | N/A | Africa | Benin | Prof. Eric Schranz |
| Zambia | TOT8933 | VI059557 | Africa | Zambia | Prof. Eric Schranz |
| Uganda-1 | TOT8889 | VI059513 | Africa | Uganda | Prof. Eric Schranz |
| Uganda-2 | TOT8887 | VI059511 | Africa | Uganda | Prof. Eric Schranz |
| Malawi | TOT8918 | VI059542 | Africa | Malawi | Prof. Eric Schranz |

<sup>1</sup> Identification codes from the AVGRIS (AVRDC's Vegetable Genetic Resources Information System) database, http://seed.worldveg.org/. <sup>2</sup> Available from B&T World Seeds, http://b-and-t-world-seeds.com/

**Supplementary Figure 2.**
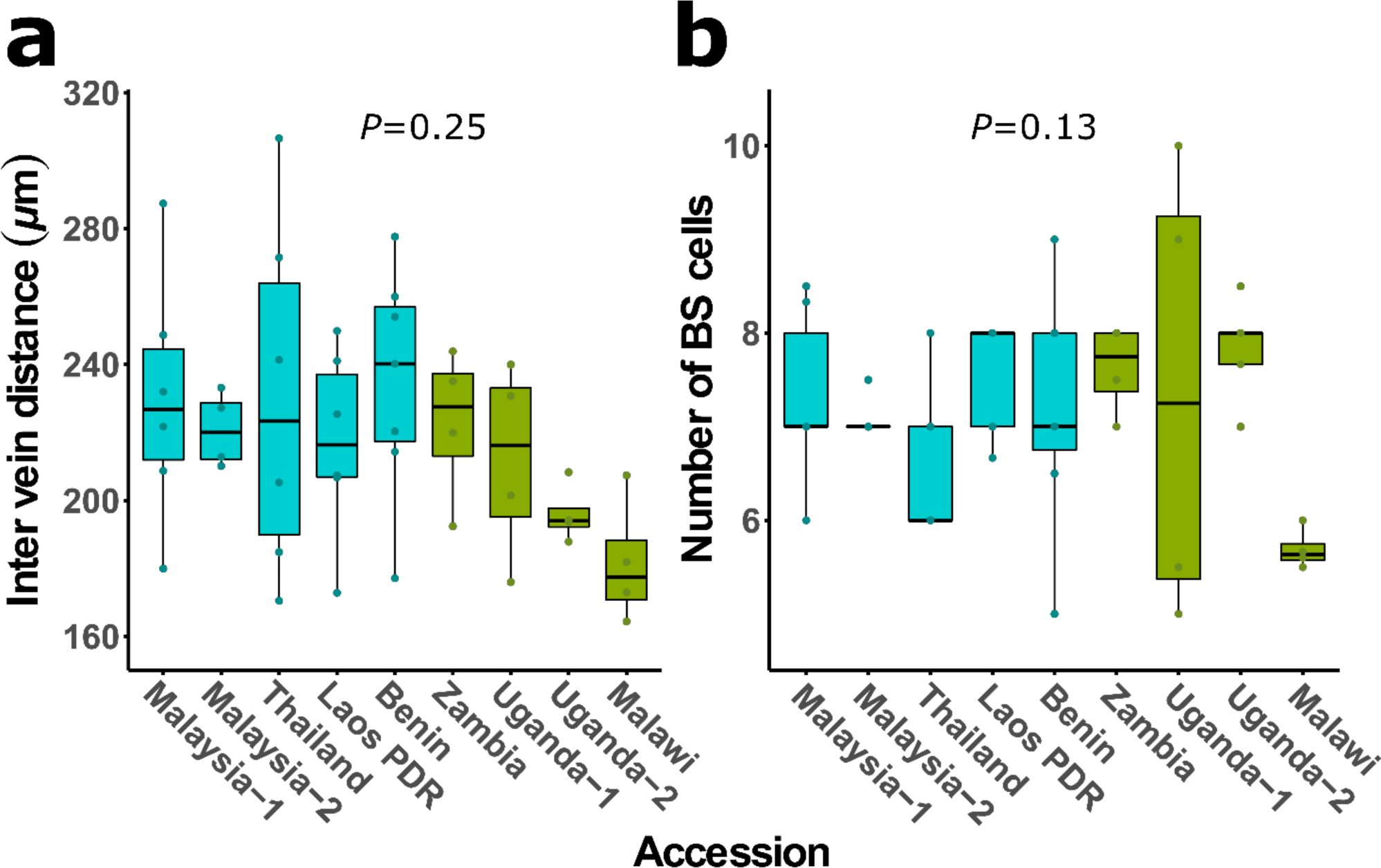
Non-variable features of Kranz anatomy among accessions. **a**, Inter-vein distance, **b**, the number of bundle sheath (BS) cells in each BS ring. *P*-value is indicate for one-way ANOVA.

**Supplementary Figure 3.**
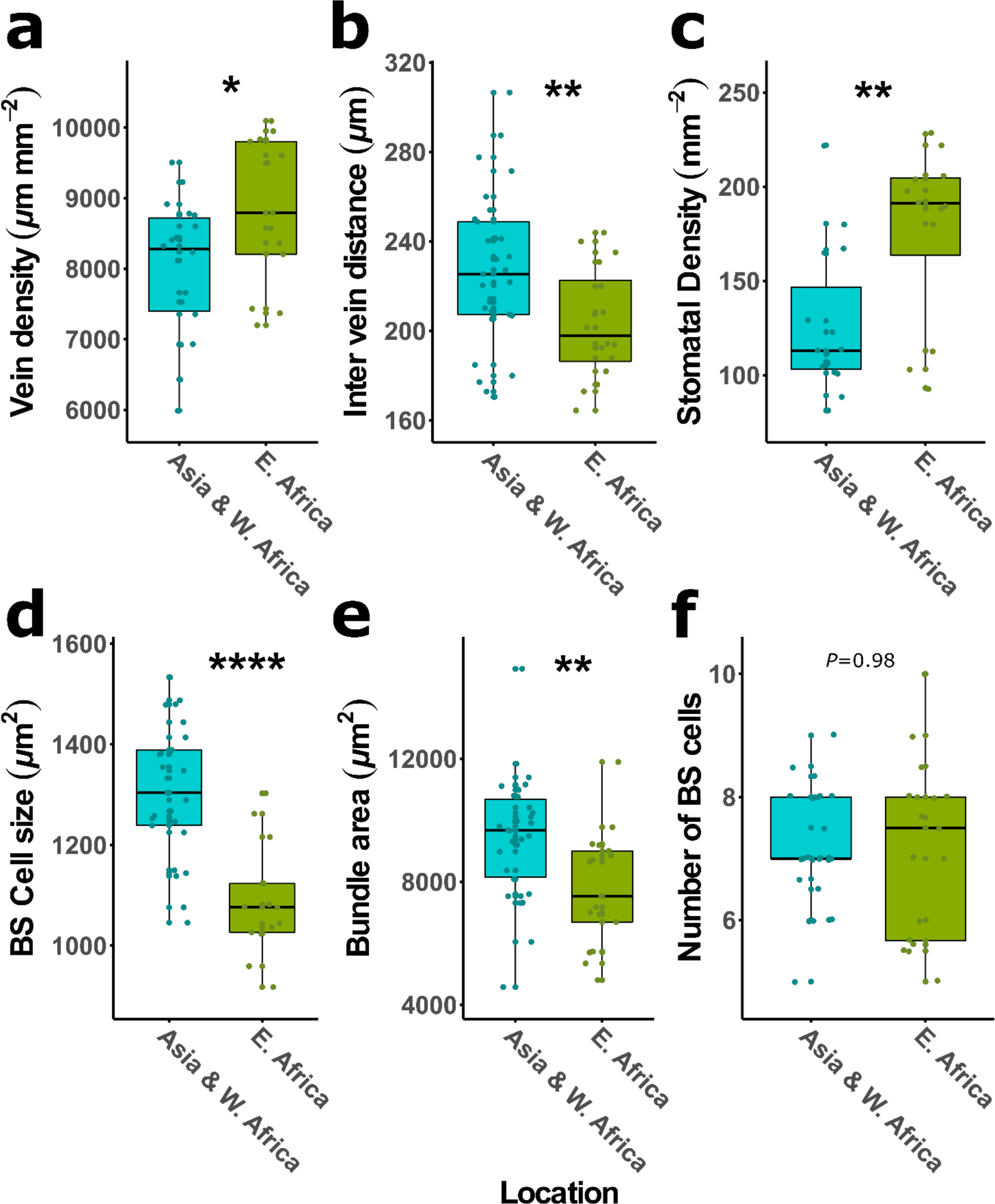
Natural variation in features in Kranz anatomy between Asian and West African accessions compared to East African accessions of *G. gynandra*. **a-f**, Vein density, inter-vein distance, average stomatal density, average bundle sheath (BS) cell size, BS area, and number of BS cells per bundle. Asterisks indicate significant differences between accessions by phylogenetic relatedness (Student’s t-test, \**P*<0.05, \*\**P*<0.01, \*\*\**P*<0.001, \*\*\*\**P*<0.0001).

**Supplementary Figure 4.**
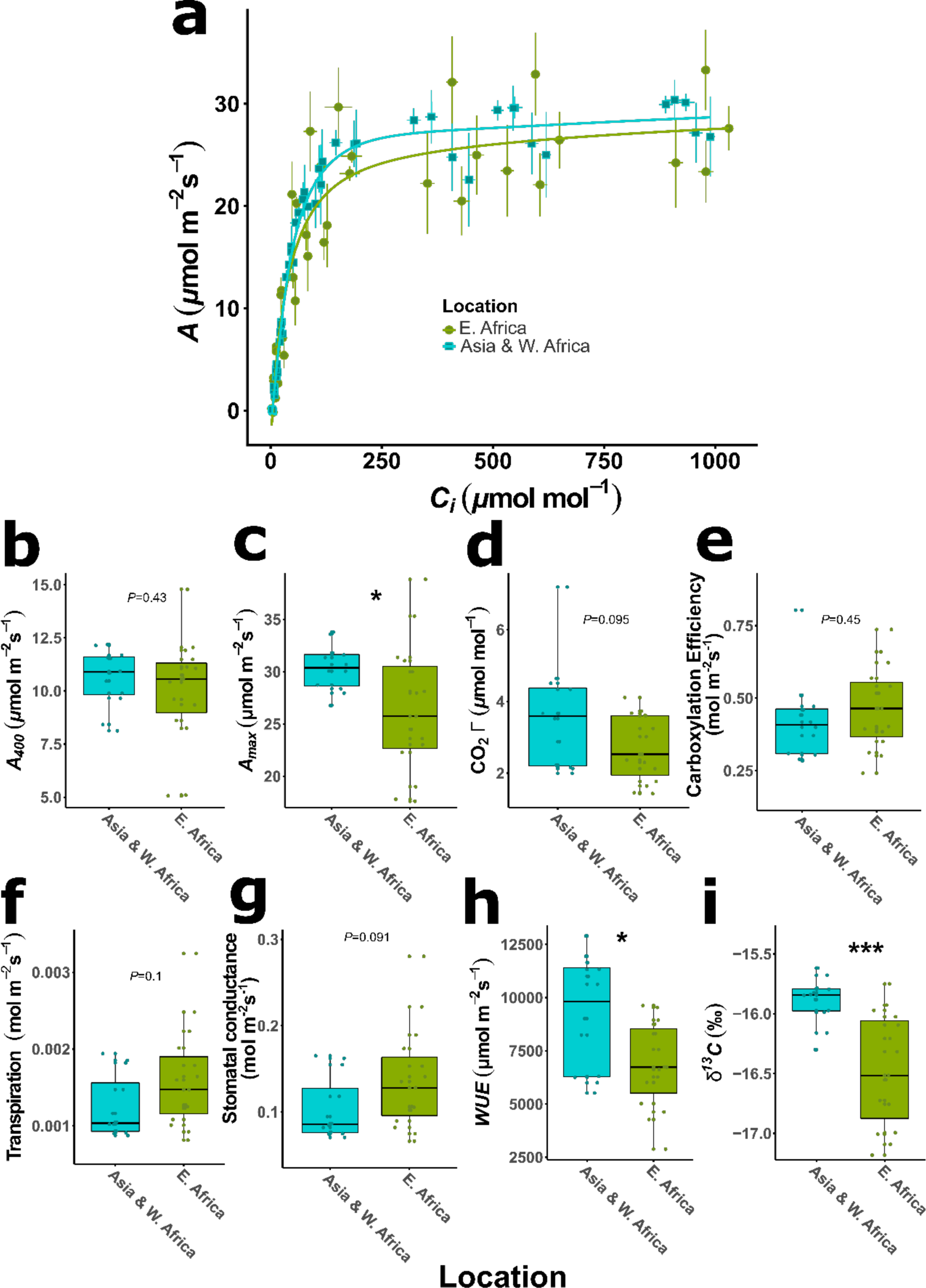
Physiological variation for photosynthetic gas exchange parameters between Asian and West African accessions compared to East African accessions of *G. gynandra*. **a**, assimilation (*A*) verses internal CO_2_ (*Ci*) curve. b-i, differences among accessions for ambient assimilation (*A*_*400*_) rates (PPFD 350 µmol m^-2^s^-1^, 400 ppm atmospheric [CO_2_], *C*_*a*_), maximal assimilation (*A*_*max*_) rates (PPFD 2000 µmol m^-2^s^-1^, 1200 ppm *C*_*a*_), CO_2_ compensation point (*Γ*), carboxylation efficiency, transpiration, stomatal conductance, water use efficiency (*WUE*), and carbon isotope composition (*δ*^*13*^*C*). Asterisks indicate significant differences between accessions by phylogenetic relatedness (Student’s t-test, \**P*<0.05, \*\**P*<0.01, \*\*\**P*<0.001, \*\*\*\**P*<0.0001, n=15 for Asia and W. Africa, n=12 for E. Africa).

**Supplementary Table 2.** Pearson product-moment correlation coefficients for Kranz anatomy traits

|  | Vein Density | Inter-vein distance | BS area | No BS cells | BS Cell size | Stomatal density |
| --- | --- | --- | --- | --- | --- | --- |
| **Vein Density** | 1.00000<br>0 | -0.67055<sup>1</sup><br>0.0001<sup>2</sup> | -0.58132<br>0.0015 | -0.05299<br>0.7929 | -0.57090<br>0.0019 | 0.56796<br>0.0020 |
| **Inter-vein distance** |  | 1.00000<br>0 | 0.70598<br><.0001 | 0.14009<br>0.4859 | 0.64649<br>0.0003 | -0.66987<br>0.0001 |
| **BS area** |  |  | 1.00000<br>0 | 0.41457<br>0.0316 | 0.80687<br><.0001 | -0.49121<br>0.0093 |
| **No BS cells** |  |  |  | 1.00000<br>0 | -0.19126<br>0.3393 | 0.01588<br>0.9373 |
| **BS Cell size** |  |  |  |  | 1.00000<br>0 | -0.50532<br>0.0072 |
| **Stomatal density** |  |  |  |  |  | 1.00000<br>0 |

<sup>1</sup>Correlation coefficient ( $\rho$ ), <sup>2</sup>P-value, under H<sub>0</sub>: $\rho=0$ ; n=27.

**Supplementary Table 3.**
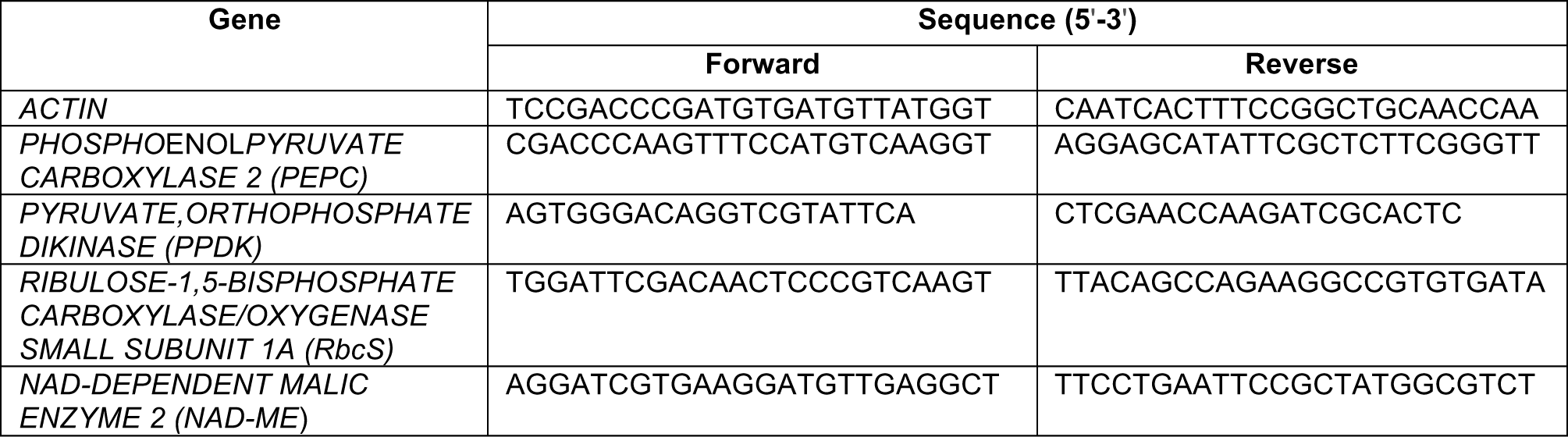
List of primers for qRT-PCR analyses

## References

1. Hibberd J. M., Sheehy J. E.& Langdale J. A. Using C4 photosynthesis to increase the yield of rice—rationale and feasibility. Curr. Opin. Plant Biol. 11, 228–231 (2008).

2. von Caemmerer S., Quick W. P.& Furbank R. T. The development of C4 rice: current progress and future challenges. Science 336, 1671–1672 (2012).

3. Hatch M. D. C4 photosynthesis: a unique elend of modified biochemistry, anatomy and ultrastructure. Biochim. Biophys. Acta - Rev. Bioenerg. 895, 81–106 (1987).

4. Kajala K. et al. Strategies for engineering a two-celled C4 photosynthetic pathway into rice. J. Exp. Bot. 62, 3001–3010 (2011).

5. Schuler M. L., Mantegazza O.& Weber A. P. M. Engineering C4 photosynthesis into C3 chassis in the synthetic biology age. Plant J. 87, 51–65 (2016).

6. Bowes G., Ogren W. L.& Hageman R. H. Phosphoglycolate production catalyzed by ribulose diphosphate carboxylase. Biochem. Biophys. Res. Commun. 45, 716–722 (1971).

7. Sharkey T. D. Estimating the rate of photorespiration in leaves. Physiol. Plant. 73, 147–152 (1988).

8. Garner D. M. G., Mure C. M., Yerramsetty P.& Berry J. O. Kranz Anatomy and the C4 Pathway. in eLS, John Wiley & Sons, Ltd, (2001).

9. Lundgren M. R., Osborne C. P.& Christin P.-A. Deconstructing Kranz anatomy to understand C4 evolution. J. Exp. Bot. 65, 3357–3369 (2014).

10. Marshall D. M. et al. Cleome, a genus closely related to Arabidopsis, contains species spanning a developmental progression from C3 to C4 photosynthesis. Plant J. 51, 886–896 (2007).

11. Sage R. F. A portrait of the C4 photosynthetic family on the 50th anniversary of its discovery: species number, evolutionary lineages, and Hall of Fame. J. Exp. Bot. (2016).

12. Aubry S., Kelly S., Kümpers B. M. C., Smith-Unna R. D.& Hibberd J. M. Deep Evolutionary Comparison of Gene Expression Identifies Parallel Recruitment of Trans-Factors in Two Independent Origins of C4 Photosynthesis. PLoS Genet 10, e1004365 (2014).

13. Lauterbach M. et al. De novo Transcriptome Assembly and Comparison of C3, C3-C4, and C4 Species of Tribe Salsoleae (Chenopodiaceae). Frontiers in Plant Science 8, 1939 (2017).

14. Bräutigam A., Schliesky S., Külahoglu C., Osborne C. P.& Weber A. P. M. Towards an integrative model of C4 photosynthetic subtypes: insights from comparative transcriptome analysis of NAD-ME, NADP-ME, and PEP-CK C4 species. J. Exp. Bot. 65, 3579–3593 (2014).

15. Reeves G., Grangé-Guermente M. J.& Hibberd J. M. Regulatory gateways for cell-specific gene expression in C4 leaves with Kranz anatomy. J. Exp. Bot. 68, 107–116 (2017).

16. Mauricio R. Mapping quantitative trait loci in plants: uses and caveats for evolutionary biology. Nat. Rev. Genet. 2, 370 (2001).

17. Oakley J. C., Sultmanis S., Stinson C. R., Sage T. L.& Sage R. F. Comparative studies of C3 and C4 Atriplex hybrids in the genomics era: physiological assessments. J. Exp. Bot. 65, 3637–3647 (2014).

18. Brown H. R.& Bouton J. H. Physiology and Genetics of Interspecific Hybrids Between Photosynthetic Types. Annu. Rev. Plant Physiol. Plant Mol. Biol. 44, 435–456 (1993).

19. Ueno O.& Sentoku N. Comparison of leaf structure and photosynthetic characteristics of C-3 and C-4 Alloteropsis semialata subspecies. Plant Cell Environ. 29, 257–268 (2006).

20. Lundgren M. R. et al. Evolutionary implications of C3 ‐C4 intermediates in the grass Alloteropsis semialata. Plant. Cell Environ. 39, 1874–1885 (2016).

21. Sogbohossou E. O. D. et al. A roadmap for breeding orphan leafy vegetable species: a case study of Gynandropsis gynandra (Cleomaceae). Hortic. Res. 5, 2 (2018).

22. Feodorova T. A., Voznesenskaya, E. V, Edwards G. E. & Roalson E. H. Biogeographic patterns of diversification and the origins of C4 in Cleome (Cleomaceae). Syst. Bot. 35, 811–826 (2010).

23. Marshall D. M. et al. Cleome, a genus closely related to Arabidopsis, contains species spanning a developmental progression from C3 to C4 photosynthesis. Plant J. 51, 886–896 (2007).

24. Brown N. J., Parsley K.& Hibberd J. M. The future of C4 research - maize, Flaveria or Cleome? Trends Plant Sci. 10, 215–221 (2005).

25. Sage R. F.& McKown A. D. Is C4 photosynthesis less phenotypically plastic than C3 photosynthesis?. J. Exp. Bot. 57, 303–317 (2006).

26. Westhoff P.& Gowik U. Evolution of C4 Photosynthesis—Looking for the Master Switch. Plant Physiol. 154, 598 LP-601 (2010).

27. Williams B. P., Johnston I. G., Covshoff S.& Hibberd J. M. Phenotypic landscape inference reveals multiple evolutionary paths to C4 photosynthesis. Elife 2, e00961 (2013).

28. Sage R. F. Tansley review: The evolution of C4 photosynthesis. New Phytol 161, 30 (2004).

29. Long S. P., Marshall-Colon A.& Zhu X.-G. Meeting the Global Food Demand of the Future by Engineering Crop Photosynthesis and Yield Potential. Cell 161, 56–66 (2015).

30. Ort D. R. et al. Redesigning photosynthesis to sustainably meet global food and bioenergy demand. Proc. Natl. Acad. Sci. 112, 8529–8536 (2015).

31. Schneider C. A., Rasband W. S.& Eliceiri K. W. NIH Image to ImageJ: 25 years of image analysis. Nat. Methods 9, 671–675 (2012).

32. Royles J. et al. Moss stable isotopes (carbon-13, oxygen-18) and testate amoebae reflect environmental inputs and microclimate along a latitudinal gradient on the Antarctic Peninsula. Oecologia 181, 931–945 (2016).

33. Levene H. in Contributions to Probability and Statistics: Essays in Honor of Harold Hotelling 278–292, Stanford University Press, (1960).

34. Duncan D. B. Multiple Range and Multiple F Tests. Biometrics 11, 1–42 (1955).

35. Pearson K. Notes on regression and inheritance in the case of two parents. Proc. R. Soc. London 58, 240–242 (1895).

36. Burgess S. J. et al. Ancestral light and chloroplast regulation form the foundations for C4 gene expression. Nat. Plants 2, 16161 (2016).

